# Localization and Quantification of Post-Translational Modifications of Proteins Using Electron Activated Dissociation Fragmentation on a Fast-Acquisition Time-of-Flight Mass Spectrometer

**DOI:** 10.1101/2023.04.29.538826

**Authors:** Joanna Bons, Christie L. Hunter, Rita Chupalov, Jason Causon, Alexandra Antonoplis, Jacob P. Rose, Brendan MacLean, Birgit Schilling

## Abstract

Protein post-translational modifications (PTMs) are crucial and dynamic players in a large variety of cellular processes and signaling, and proteomic technologies have emerged as the method of choice to profile PTMs. However, these analyses remain challenging due to potential low PTM stoichiometry, the presence of multiple PTMs per proteolytic peptide, PTM site localization of isobaric peptides, and labile PTM groups that lead to neutral losses. Collision-induced dissociation (CID) is commonly used for to characterize PTMs, but the application of collision energy can lead to neutral losses and incomplete peptide sequencing for labile PTM groups. In this study, we compared CID to an alternative fragmentation, electron activated dissociation (EAD), operated on a recently introduced fast-acquisition quadrupole-time-of-flight (QqTOF) mass spectrometer. We analyzed a series of synthetic modified peptides, featuring phosphorylated, succinylated, malonylated, and acetylated peptides. We performed targeted, quantitative parallel reaction monitoring (PRM or MRM^HR^) assays to assess the performances of EAD to characterize, site-localize and quantify peptides with labile modifications. The tunable EAD kinetic energy allowed the preservation of labile modifications and provided better peptide sequence coverage with strong PTM-site localization fragment ions. Zeno trap activation provided significant MS/MS sensitivity gains by an average of 6–11-fold for EAD analyses, regardless of modification type. Evaluation of the quantitative EAD PRM workflows revealed high reproducibility with coefficients of variation of typically ∼2%, as well as very good linearity and quantification accuracy. This novel workflow, combining EAD and Zeno trap, offers confident, accurate, and robust characterization and quantification of PTMs.

## INTRODUCTION

The proteome of any organism provides an impressive complexity that tailors and fine-tunes protein functions, signaling, and interactions. Each protein can exhibit different and multiple modifications, different splice variants and even protein proteolysis and degradation products. Proteome complexity and dynamic changes are closely associated with protein post-translational modifications (PTMs), as multiple protein isoforms, also referred to as proteoforms, can result from a single RNA transcript. Indeed, the reversible or irreversible covalent addition of a chemical group or moiety to an amino acid residue within a protein is typically completed via fast kinetic reactions, either enzymatically or chemically, and often changes the physiochemical properties of a protein. As a consequence, PTMs elegantly regulate protein function, conformation, protein-protein interactions, and even organelle localization and, thus, play pivotal roles in a multitude of cellular and epigenetic processes and signaling.

Hundreds to thousands of different PTMs have been investigated. They are chemically diverse and occur on many different protein amino acid residues ^1^. For instance, acylation, including acetylation ^2, 3^, succinylation ^4, 5^ and malonylation ^6^, modifies lysine residues, thereby neutralizing the positive charge of lysine (e.g., acetylation) ^7^, or even changing the net charge of lysine from a positive (+1) to a negative (-1) charge (e.g., succinylation and malonylation) ^8^ at physiological pH. Significant research efforts have been dedicated to protein phosphorylation due to its relevance in protein signaling and signaling cascades, activation of kinases via (auto)phosphorylation, and the highly dynamic interplay between kinases and phosphatases to add or remove the modifications. Phosphorylation primarily modifies serine residues, followed by threonine and tyrosine residues, in eukaryotes, but can also be present on histidine and aspartic acid residues in prokaryotes, and these modifications can lead to strong structural rearrangements ^9^. Changes in PTM profiles are associated with dysregulated biological mechanisms and diseases, such as metabolism dysfunction, inflammation, neurodegenerative diseases, cancer, and cardiovascular disease ^10–12^. Highly specific kinase inhibitors, for example, provide popular therapeutic intervention mechanisms for diseases ^13–16^.

Proteomic workflows using liquid chromatography–tandem mass spectrometry (LC-MS/MS) were established as powerful tools for the proteome-wide profiling of PTMs and to discover novel modifications, as these changes induce distinct mass shifts detectable in the MS/MS fragment ion spectra (e.g., +42.01 m/z for acetyl, +86.00 m/z for malonyl, +100.02 m/z for succinyl, and +79.97 m/z for phosphoryl) ^17, 18^. However, analytic workflows with LC-MS/MS for PTM analysis still encounter many challenges, such as neutral losses from labile modifications during MS/MS, low PTM stoichiometry resulting in low abundance of the modified peptides, post-translationally modified peptide isomers, and multiple modifications per proteolytic peptide. These can lead to incorrect PTM identification, ambiguous site-localization and inaccurate quantification, especially as most analyses rely on the quality of only one MS/MS spectrum per PTM site and the detection of PTM-containing fragment ions to determine PTM presence and localization. Collision-induced dissociation (CID) is commonly used to characterize PTMs. However, because collisional activation leads to cleavage of the weakest bonds (typically amino acid amide bonds of the peptide backbone) ^19^, CID can cause neutral losses from the labile PTM groups ^6, 20^. Alternatively, electron-based dissociation methods, such as electron capture dissociation (ECD) ^21^ and electron transfer dissociation (ETD) ^22^, were developed to cope with these limitations by maintaining the labile PTM group intact and generating more complete fragment ion series ^23, 24^. However, the latter methods often suffer from low fragmentation efficiency of the precursor ions, specifically for doubly charged precursor ions ^23, 25^ that are subsequently not selected for fragmentation. MS/MS fragmentation via ECD and ETD primarily selects higher charge states of the precursor ions ^19, 26^; however, those require relatively long reaction times to achieve optimal fragmentation ^24, 27^. In 2015, Baba *et al.* introduced a high-throughput ECD device coupled to an orthogonal quadrupole-time-of-flight (QqTOF) mass spectrometer using a novel branched radio frequency ion trap, that is capable of performing ECD fragmentation with enhanced fragmentation frequency and shorter trapping periods ^28^.

Most recently, in 2021, an electron activated dissociation (EAD) device with redesigned electron beam optics was implemented on a novel fast scanning QqTOF mass spectrometer ^29^. Briefly, the optimized design increases the electron beam intensity for efficient and fast EAD. With this device design, EAD experiments can be performed at reaction times as fast as 10 msec with a range of electron kinetic energies (KEs) (0–15 eV) for all positive precursor ion charge states. In addition, EAD is compatible with fast high-performance liquid chromatography (HPLC) separations using short gradients for LC-MS/MS acquisitions. Strikingly, easily tunable electron KEs enable EAD experiments to be performed with different distinctly controlled energy inputs on singly and multiply charged analytes. EAD mechanisms include low-energy ECD and higher-energy “hot ECD” ^30^, which relies on irradiating analytes with electrons with higher KEs of several electron volts.

To maximize duty cycle efficiency and to achieve higher sensitivity, the QqTOF mass spectrometer also integrates a novel Zeno trap technology ^31^. The Zeno trap device is located in between the Q2 quadrupole collision cell and the TOF accelerator. It efficiently minimizes the duty cycle ion loss when ions are released into the TOF accelerator. Briefly, ions are captured and stored in the Zeno trap after exiting the collision cell. Then, in synchronization with the TOF accelerator, the ions are released, based on potential energy, from higher m/z ions to lower m/z ions, to simultaneously and spatially focus all ions at the center of the accelerator plate at the same time for efficient delivery into the TOF region for detection ^31^. This technology improves the ion duty cycle to >90% through this region of the MS system and, thus, enhances detection sensitivity, which represents an asset for measurements of EAD and low-abundance PTM peptides.

In this study, we assessed the performances of EAD and CID to confidently and accurately identify, site localize and quantify PTMs. We analyzed a series of biologically relevant synthetic modified peptides, featuring malonylation, succinylation, acetylation, and phosphorylation, using targeted parallel reaction monitoring (PRM or MRM^HR^) assays ^32^ performed on a QqTOF system (**Figure 1**). We explored a range of EAD KE values to determine optimal values, which keep labile modifications intact, and provide comprehensive peptide fragmentation coverage, generating strong PTM site-localizing ions, with particular focus on malonylation and phosphorylation. We also investigated the capabilities of Zeno trap activation to increase sensitivity when combined to EAD fragmentation.

**Figure 1.**
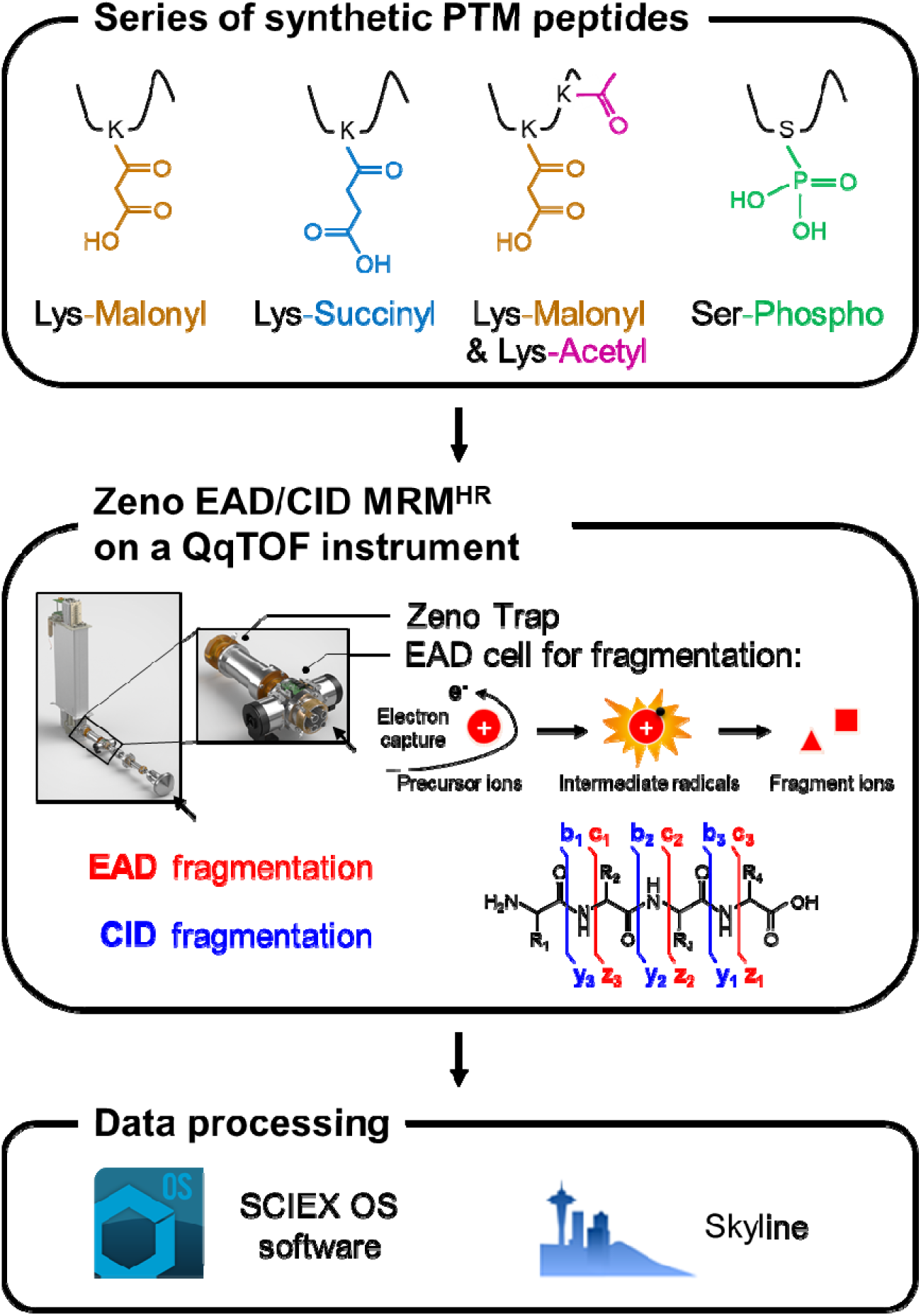
Study design for efficient PTM analysis. *A series of synthetic modified peptides, featuring lysine malonylation, lysine succinylation, doubly lysine malonylation and lysine acetylation, and serine phosphorylation, were analyzed using microLC on a fast-scanning quadrupole-time-of-flight (QqTOF) mass spectrometer using targeted analysis assays (termed PRM or MRM^HR^). Collision-induced dissociation (CID) or electron activated dissociation (EAD) was applied, and the effects of Zeno trap activation were also assessed. The EAD reaction was performed in the EAD cell located upfront of the collision cell/quadrupole 2 (Q2), and the Zeno trap device was directly behind Q2. Finally, the Zeno EAD PRM and Zeno CID PRM data were analyzed with SCIEX OS software and Skyline to detect, visualize and quantify c and radical z ions generated by EAD and c and y ions by CID.*

## METHODS

### Synthetic peptides

Synthetic malonylated, succinylated, acetylated, doubly malonylated-acetylated, and phosphorylated peptides are detailed in **Table S1**. The malonylated synthetic peptide TVDGPSG**Kma**LWR* containing a stable isotope-labeled arginine residue (R* with ^13^C_6_ and ^15^N_4_) was obtained from Thermo Fisher Scientific at >90% chemical purity and >99% isotopic purity. The phosphorylated synthetic peptides LITVDGNIC**pS**GKSK and LITVDGNICSGK**pS**K were from Princeton Biomolecules Corporation at >99% chemical purity. Succinylated synthetic peptides containing a stable isotope-labeled arginine (^13^C_6_, ^15^N_4_), phenylalanine (^13^C_9_, ^15^N_1_), or valine (^13^C_5_, ^15^N_1_) residue were from New England Peptide at >99% chemical and >99% isotopic purity, and additional phosphorylated and doubly malonylated-acetylated synthetic peptides were from Vivitide at >95% chemical purity (99% isotopic purity). Synthetic peptides were diluted in 2% acetonitrile, 0.1% formic acid in H_2_O for MS analysis.

### LC-MS/MS analysis

LC-MS/MS analyses were conducted on either a nanoLC 425 system (SCIEX, Concord, CA) or Waters UHPLC system (Waters, Milford, MA), coupled to a ZenoTOF 7600 mass spectrometer (SCIEX; referred to as QqTOF throughout) and an OptiFlow Turbo V ion source with a microflow probe and a 25-μm electrode. The solvent system consisted of 100% water with 0.1% formic acid (solvent A) and 100% acetonitrile with 0.1% formic acid (solvent B). Peptides were directly loaded and eluted at 5 μL/min on a 0.3 mm x 150 mm (2.6-μm particle size) Phenomenex Omega Polar analytical column, using the following gradient of solvent B: peptides were loaded with 3% solvent B, and separated using either a 5-, 8-, or 13-min linear gradient from 3 to 35% solvent B, followed by an increase to 80% solvent B for 0.5 min, a hold at 80% solvent B for 2 min, a decrease to 3% solvent B over 0.5 min, and a hold at 3% solvent B for 4 min. The total HPLC acquisition lengths were 12, 15 and 20 min, respectively, depending on the different experimental designs. Data were collected in PRM/MRM^HR^ mode as detailed in **Table S2**. For CID fragmentation, collision energy values were determined for each peptide based on their charge state and m/z using predetermined collision energy equations available for the ZenoTOF 7600 system (QqTOF). For EAD fragmentation, kinetic energies were ramped from 0 eV to 11 eV and optimized for each analyte. Electron current was also ramped and a final optimized value of 5,000 nA was used throughout the study. An EAD reaction time of 10 msec was used in all experiments.

### Data processing

PRM data were processed using SCIEX OS software 2.1 using both Explorer and Analytics modules, as well as Skyline-daily (version 21.2.1.455) ^33^. The Explorer module in the SCIEX OS software was used to visualize the annotated spectra and the Analytics module to integrate peak areas, to assess EAD kinetic energy ramping data, Zeno trap ON/OFF sensitivity gains, and the EAD PRM quantification accuracy and reproducibility. More particularly, the workflow “Quantitation and targeted identification” was selected, “MQ4” Integration Algorithm and “Relative Noise” Signal-to-Noise Algorithm were used, and default settings were applied, except that XIC width was set at 0.05 Da. Extracted ion chromatograms (XICs) were manually verified, and the integration was refined if necessary. Skyline-daily (version 21.2.1.455) was used to analyze the phosphorylated synthetic peptides LITVDGNIC**pS**GKSK and LITVDGNICSGK**pS**K from EAD PRM and CID PRM experiments. Briefly, product ions (singly and doubly charged y-, b-, z-, and c-type ions) from “ion 1” to “last ion” per precursor ion were extracted. All matching scans were used. Skyline-daily (version 21.2.1.455) was also used to analyze the succinylated peptides, investigating EAD PRM quantification linearity. Here, product ions (singly to triply charged z.-, z′-, and c-type ions) from ion 1 to last ion per precursor ion were extracted. All matching scans were used. XICs were manually verified and integrated.

### Data availability

Raw data have been uploaded to the MassIVE repository of the Center for Computational Mass Spectrometry at UCSD, and can be downloaded using the following link: https://massive.ucsd.edu/ProteoSAFe/dataset.jsp?task=d1ee2225612544c4bfd77141bd ddb021 (MassIVE ID number: MSV000091722).

[Note to the reviewers: To access the data repository MassIVE (UCSD) for MS data, please use: Username: MSV000091722_reviewer; Password: winter].

## RESULTS AND DISCUSSION

To investigate the performance of the newly introduced platform for electron activated dissociation (EAD) fragmentation to identify, localize and quantify a diverse set of PTMs, a series of synthetic malonylated, succinylated, doubly malonylated-acetylated and phosphorylated peptides (**Table S1**) were analyzed on a fast-scanning QqTOF mass spectrometer using targeted EAD PRM/MRM^HR^ and CID PRM/MRM^HR^ approaches (**Figure 1**) with and without Zeno trap activation. EAD provides reagent-free electron-based fragmentation methods, operated within the EAD device, which contains a cathode for generating the electron beam, and which is located upstream of the quadrupole collision cell on the instrument. Briefly, trapped multiply protonated precursor ions interact with electrons, producing intermediate radical species. Importantly, for EAD, the kinetic energy (KE) of the electron beam can be tuned from 0 to 25 eV to obtain optimized conditions for different analytes. These intermediate radical species undergo further fragmentation; the N-C_α_ bond is cleaved, and C-terminal radical z ions, or z+1 ions, and N-terminal c ions are produced. Finally, EAD PRM and CID PRM data were analyzed with the SCIEX OS software and with the open-source tool Skyline ^33^, that now supports z+1 radical ion (z^•^ ion) and z+2 ion (z’ ion) type (implemented in Skyline version 21.2.1.404).

### Preservation of labile modifications and prevention of neutral losses upon fragmentation using EAD

To explore the capabilities of EAD to preserve labile modifications, the malonylated tryptic peptide TVDGPSG**^192^Kmal**LWR derived from mouse glyceraldehyde-3-phosphate dehydrogenase (P16858) was synthetized with a malonylated lysine at position K-192 and analyzed using EAD PRM (**Figure 2A**) and CID PRM (**Figure 2B**). EAD fragmentation generated intact z+1 and c fragment ions (**Figure 2A**) and a near complete sequence characterization (**Figure 2C**). Specifically, fragment ions ranging from z_4_+1 to z_11_+1, as well as c_9_ and c_10_, contained the labile malonyl group, and importantly, these ions did not undergo significant neutral losses using EAD, hence enabling confident PTM site localization. However, CID fragmentation mode caused significant neutral loss of CO_2_ (-44 m/z), indicative of decarboxylation from the labile malonyl group ^6^, that was observed for the entire y-ion series (**Figure 2B and D**). More specifically, the neutral loss produced ions with m/z that are indicative of an acetyl group at position K-192, subsequently leading to misidentification of the PTM with CID (m/z malonyl minus CO_2_ corresponds to m/z acetyl). Thus, EAD preserves labile PTMs, which enables confident PTM identification and provides direct evidence for PTM site localization.

**Figure 2.**
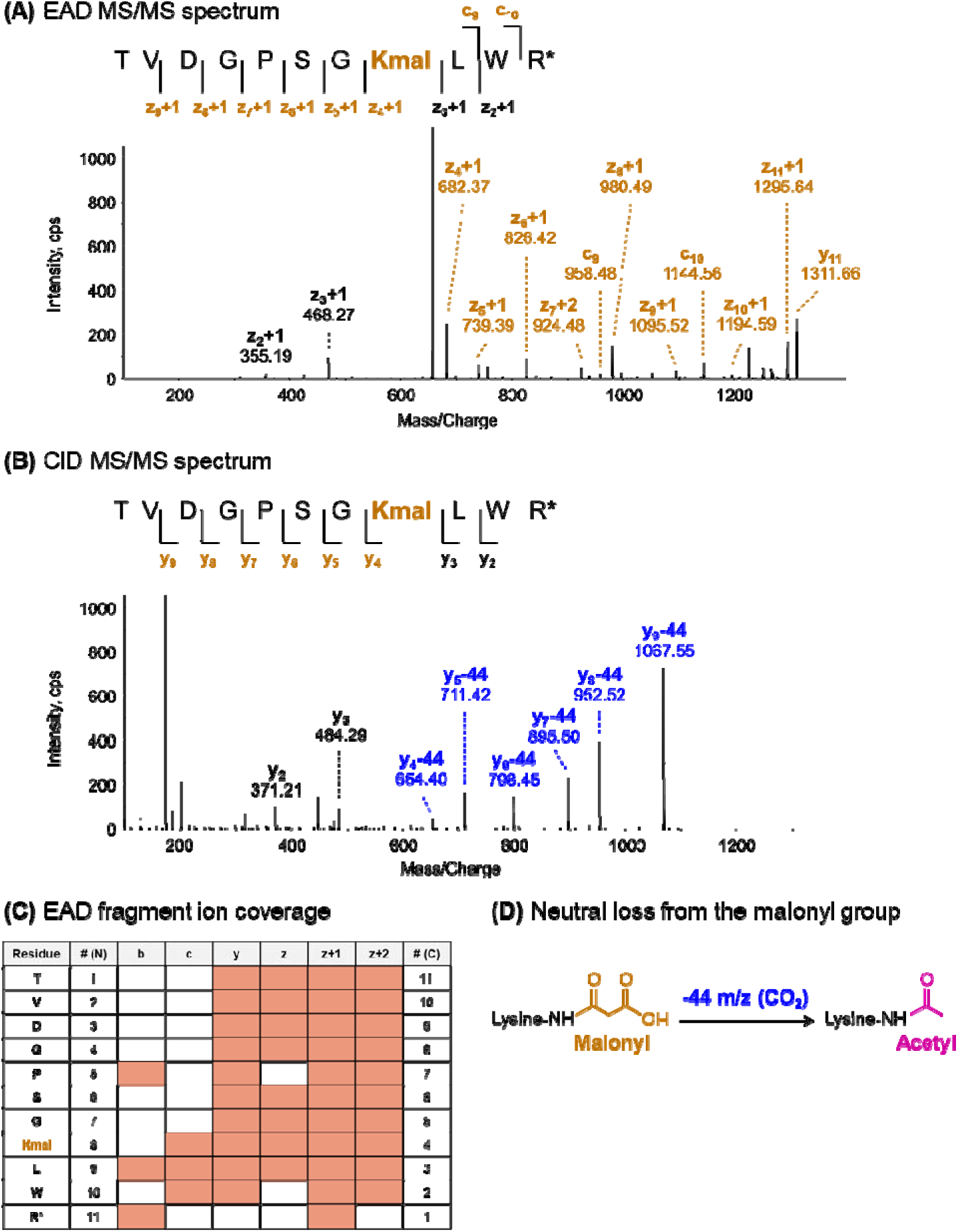
Preservation of intact labile malonyl groups with EAD. *The malonylated peptide TVDGPSG*^192^***Kmal****LWR* (precursor ion at m/z 656.33, z = 2+) from mouse glyceraldehyde-3-phosphate dehydrogenase (P16858) was analyzed using **(A)** EAD PRM at KE = 5 eV and **(B)** CID PRM. Site-specific fragment ions containing the malonyl-group are indicated in brown. **(C)** Near complete sequence characterization was achieved with the EAD dissociation. Comprehensive EAD fragmentation generated high-intensity fragment ions that provided direct evidence for PTM site localization, whereas no “intact” PTM-specific differentiating ions were detected with CID fragmentation **(D)** as a result of neutral loss of -CO_2_ (-44 m/z)*.

### Kinetic energy ramping and optimization of EAD fragmentation

One strength of ECD, including EAD, that differs from ETD dissociation is the ability to tune the kinetic energy of the electron beam specifically to the analytes investigated. To determine the optimal fragmentation parameters both to preserve labile PTM groups on modified peptides and to achieve highest sensitivity, EAD fragmentation of the peptide TVDGPSG^192^**Kmal**LWR was performed with KE values, ranging from 0 eV to 11 eV (**Figure 3**). In this study, a KE value of 5 eV generated high intensity, intact PTM site-specific fragment ions with very little background noise (**Figure 3A**). When the KE is increased to 8 eV or higher, the Zeno EAD MS/MS spectra became more complex, and neutral losses of CO_2_ (-44 m/z) from fragment ions containing the malonyl group were noted, as highlighted for z_5_+1, z_6_+1, and z_7_+1 ions (**Figure 3B**).

**Figure 3.**
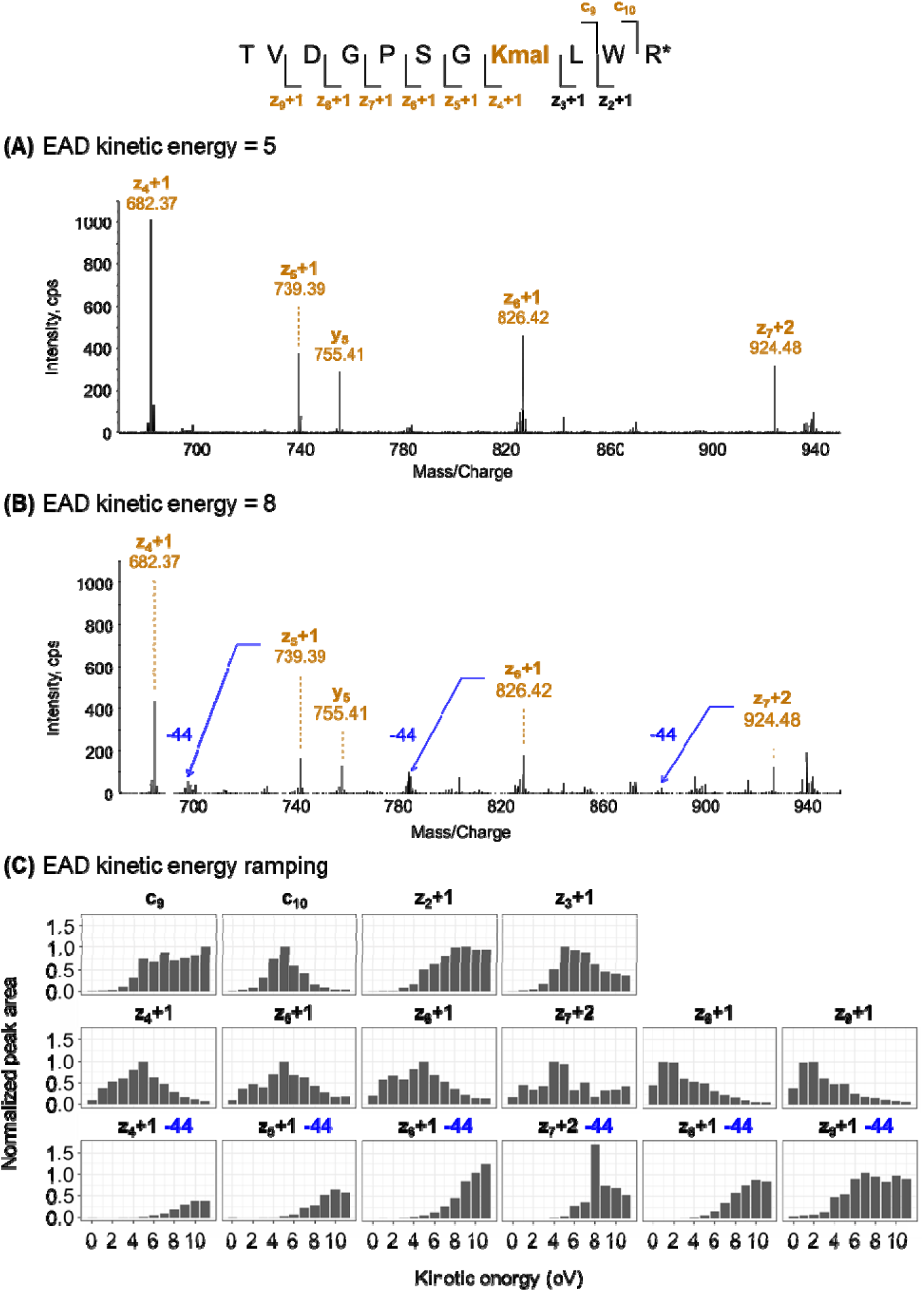
Effects of tuning EAD kinetic energy on the fragmentation pattern. *Zeno EAD MS/MS spectra of the malonylated TVDGPSG*^192^***Kmal****LWR* precursor ion (m/z 656.33, z = 2+) resulting from kinetic energy (KE) values of **(A)** 5 eV and **(B)** 8 eV. Site-specific fragment ions containing the malonyl group are indicated in brown. Displayed MS/MS are zoomed over m/z 670–950 range. **(C)** The EAD KE was ramped from 0 to 11 eV. Chromatographic peaks were extracted for 16 fragment ions. For each intact fragment ion, peak area values were normalized to the respective highest area value. For the fragment ions with a neutral loss, area values were normalized to the value of the intact ion counterpart, respective of the KE. Neutral losses of -CO_2_ (-44 m/z) were observed for higher KEs*.

As a second step, dissociation patterns were investigated for each individual fragment ions across the ramping EAD KE values (**Figure 3C**). For intact fragment ions, the peak area of the extracted ion chromatogram (XIC) was normalized to the highest peak area value obtained among the 12 KE values tested. For fragment ions with neutral loss (-44 m/z), the peak area was normalized against the area of its intact counterpart ion for each KE step, respectively. This analysis revealed that different dissociation patterns were obtained for the PTM site-specific fragment ions of TVDGPSG^192^**Kmal**LWR by EAD fragmentation. Indeed, the peak area for most fragment ions containing the labile malonyl group, namely c_10_, z_3_+1, z_4_+1, z_5_+1, and z_6_+1, was highest at KE = 5 eV. With higher KE values, some neutral PTM losses were observed, and peak areas of ions with neutral loss were at least half as intense as for their respective intact ions from KE = 8 eV (except for z_4_+1). Thus, in this study, KE = 5 eV maintained the integrity of the labile malonyl group and maximized sensitivity of site-specific fragment ions for TVDGPSG^192^**Kmal**LWR.

### Differentiation and accurate quantification of phosphopeptide isomers using EAD

The challenges in analyzing PTMs include differentiating isomers and confidently assigning the exact location of a PTM on the peptide. To determine how EAD fares in these analyses, two isomeric phosphorylated peptides from bovine mitochondrial NDUFA10 subunit of Complex I (P34942), LITVDGNIC**^56^pS**GKSK and LITVDGNICSGK**^59^pS**K, were synthetized with a phospho-serine at positions S-56 and S-59, respectively (**Figure 4**). Here, the modification sites of the two isomers are two amino acids apart and close to the peptide C-terminus; consequently, only a limited subgroup of PTM-containing fragment ions enables differentiation of the PTM isomers and localization of the PTM sites. Using CID, small y-ions and large b-ions are often difficult to detect, thus making it very difficult to characterize PTMs near the peptide C-terminus. Also, this fragmentation type can lead to neutral loss of H_3_PO_4_ (-98 m/z) from the labile phosphoryl group on serine residues ^24^ (**Figures S1A and S2A**). In contrast, Zeno EAD PRM, when performed at optimized KE = 2 eV for both phosphopeptides (**Figures S1B and S2B**), created high intensity and intact differentiating, PTM-containing fragment ions (**Figure 4A-B**). Specifically, fragment ions c_10_, c_11_ and c_12_ provided direct evidence for the phosphorylation site localization for LITVDGNIC**^56^pS**GKSK, and fragment ions z_2_+1 to z_4_+1 for LITVDGNICSGK**^59^pS**K (in blue on the Zeno EAD MS/MS spectra). Additional differentiating, non-PTM containing ions were detected, such as z_2_+1, z_3_+1 and z_4_+1 for LITVDGNIC**^56^pS**GKSK and c_10_ to c_12_ for LITVDGNICSGK**^59^pS**K (in black on the Zeno EAD MS/MS spectra). By taking advantage of the recently implemented features in Skyline to analyze both the z+1-ion and z+2-ion types from electron-based dissociation data, these six differentiating ions were extracted from the Zeno EAD PRM and Zeno CID PRM data for the two phosphopeptide isomers (**Figure 4C-F**). This revealed that very strong PTM-containing fragment ions could be detected using EAD, although their abundance was very low with CID.

**Figure 4.**
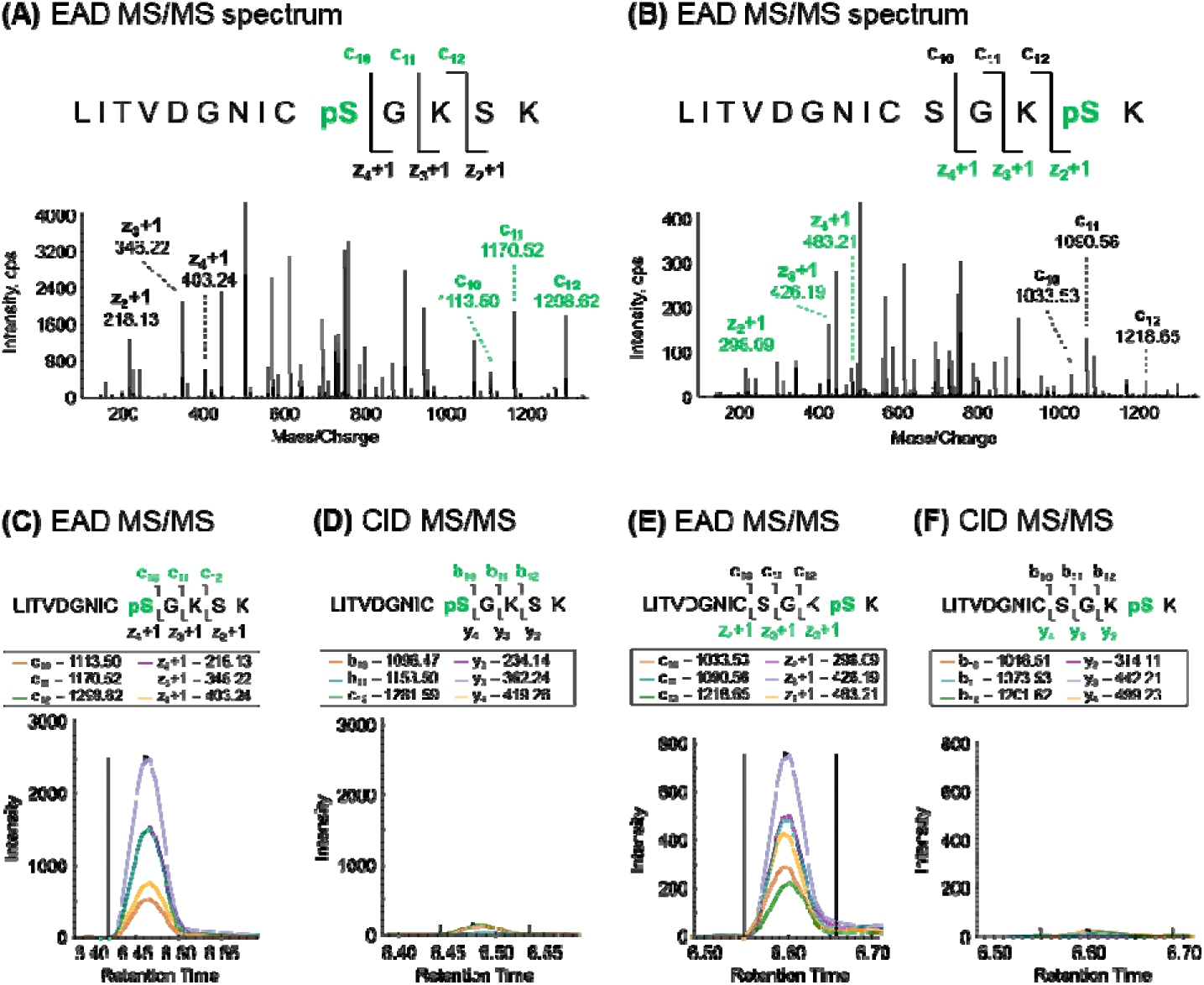
Confident differentiation of phosphorylated peptide isomers using EAD. *Zeno EAD MS/MS spectra of the phosphorylated **(A)** LITVDGNIC**^56^pS**GKSK and **(B)** LITVDGNICSGK**^59^pS**K isomer precursor ions (m/z 505.58, z = 3+) from the bovine mitochondrial NDUFA10 subunit of Complex I (P34942) are shown. EAD KE = 2 eV was applied. Site-specific fragment ions containing the phosphoryl group are indicated in green, and additional differentiating ions in black. Differentiating ions of phosphorylated isomers **(C, D)** LITVDGNIC**^56^pS**GKSK and **(E, F)** LITVDGNICSGK**^59^pS**K analyzed in **(C, E)** EAD (KE = 2 eV) and **(D, F)** CID modes were extracted in Skyline. With EAD, the labile phosphoryl group was preserved*.

To better reflect the challenge in biology of determining the actual site location of the phosphorylation when there are multiple possible sites, both phosphopeptide isomers were mixed together before Zeno EAD PRM analysis (**Figure 5**). Strikingly, near-complete c ion and z+1 ion series were detected for the two peptides. When extracting in Skyline all possible fragment ions (*i.e.*, the differentiating ions and the ions common to both isomers), two very similar, closely eluting chromatographic peaks were observed as they correspond to each of the isomers (**Figure 5A-B**). However, restricting the extraction to only the PTM site-specific ions (c_10_, c_11_, c_12_, z_2_+1, z_3_+1, z_4_+1), we were able to unambiguously differentiate these isomers, with LITVDGNIC**^56^pS**GKSK eluting at 6.45 min and LITVDGNICSGK**^59^pS**K eluting at 6.60 min (**Figure 5C-D**).

**Figure 5.**
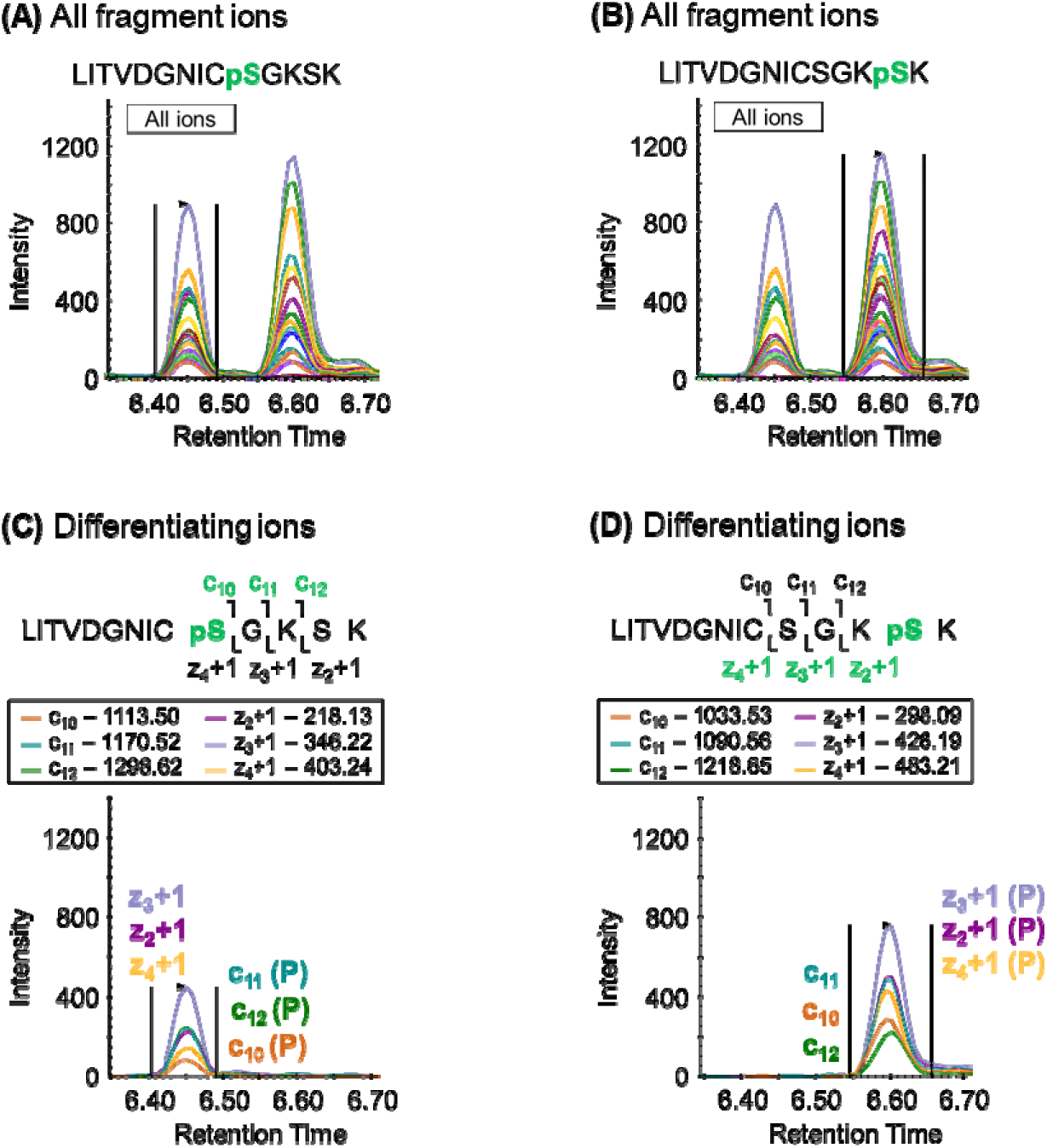
PTM site localization and quantification from EAD data using Skyline. *Near complete c- and z+1-ion series were obtained by Zeno EAD PRM acquisition for the phosphorylated isomers **(A)** LITVDGNIC**^56^pS**GKSK and **(B)** LITVDGNICSGK**^59^pS**K as shown in* Figure 4. *Post-acquisition data refinement in Skyline allowed us to differentiate the two isomeric peptides **(C)** LITVDGNIC**^56^pS**GKSK and **(D)** LITVDGNICSGK**^59^pS**K, using only discriminating ions, and to provide with quantification capabilities*.

Altogether, these results demonstrate that EAD confidently differentiates positional isomers, providing definitive and precise localization of the PTM site with sensitivity performances compatible with quantification. In addition, Skyline offers the possibility to visualize, refine, and perform precise quantification of PTMs from EAD PRM data.

### Improvement of EAD MS/MS sensitivity using the Zeno trap technology

The QqTOF instrument is equipped with the novel Zeno trap technology ^31^ that improves the duty cycle by better controlling ion transfer to the TOF accelerator. **Figure 6** illustrates the effect of Zeno trap pulsing on EAD MS/MS of the malonylated peptide TVDGPSG^192^**Kmal**LWR. Comparison of EAD MS/MS spectra acquired with and without Zeno trap activated showed significant enhancement in sensitivity provided by this feature and improved detection of the PTM site-specific fragment ions (**Figure 6A-B**). To quantify this gain, the ratio of the fragment ion XIC areas obtained with and without Zeno trap activated was determined for PTM-containing ions (**Figure 6C**). The sensitivity gain using the Zeno trap ranged from 4.0- and 8.0-fold with an average sensitivity gain of 6.1. This demonstrates the benefit of Zeno trap activation for analyzing very low abundant modified peptide ions. Similar results were obtained for the two phosphorylated peptide isomers, LITVDGNIC**^56^pS**GKSK (**Figure 7**) and LITVDGNICSGK**^59^pS**K (**Figure S3**). The combination of EAD with Zeno trap activated provided higher sensitivity with better detection of the differentiating ions (**Figure 7A-B** and **Figure S3A-B**) and an average sensitivity gain of 11.3-fold for LITVDGNIC**^56^pS**GKSK and 11.4-fold LITVDGNICSGK**^59^pS**K, with specific ranges of 7.4–14.9 and 6.8–19.9, respectively (**Figure 7C** and **Figure S3C**).

**Figure 6.**
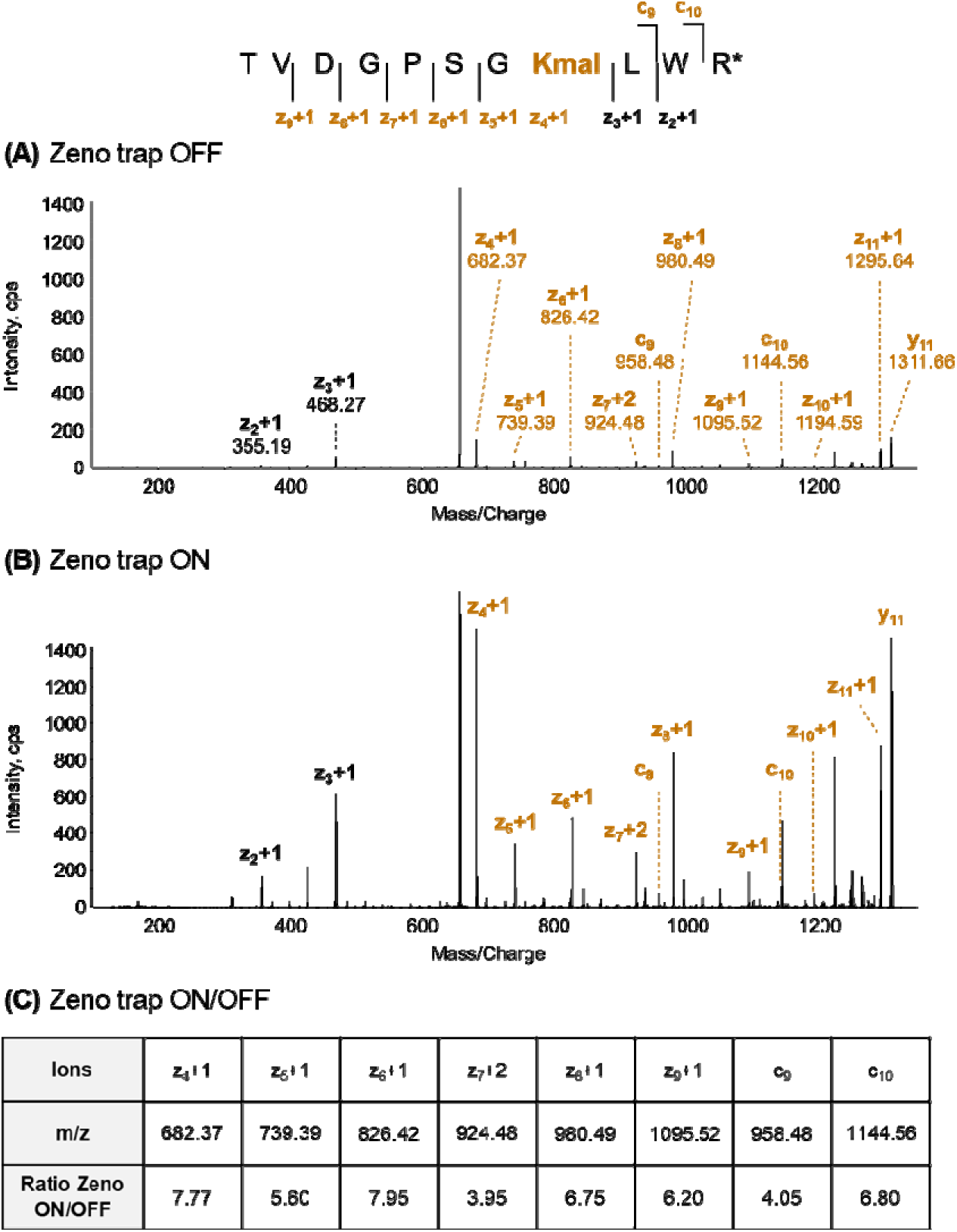
Increased EAD MS/MS sensitivity for malonylation analysis with Zeno trap technology. *EAD MS/MS spectra of the malonylated TVDGPSG*^192^***Kmal****LWR* precursor ion (m/z 656.33, z = 2+) **(A)** without and **(B)** with Zeno trap activated are shown (525 pg on-column injected). **(C)** TVDGPSG*^192^***Kmal****LWR peptide was injected at 105 pg on-column for EAD PRM analysis with and without using the Zeno trap. Chromatographic peaks were extracted, and sensitivity changes between Zeno trap on and off were determined. * Weak signal without using Zeno trap. EAD KE = 5 eV was applied*.

**Figure 7.**
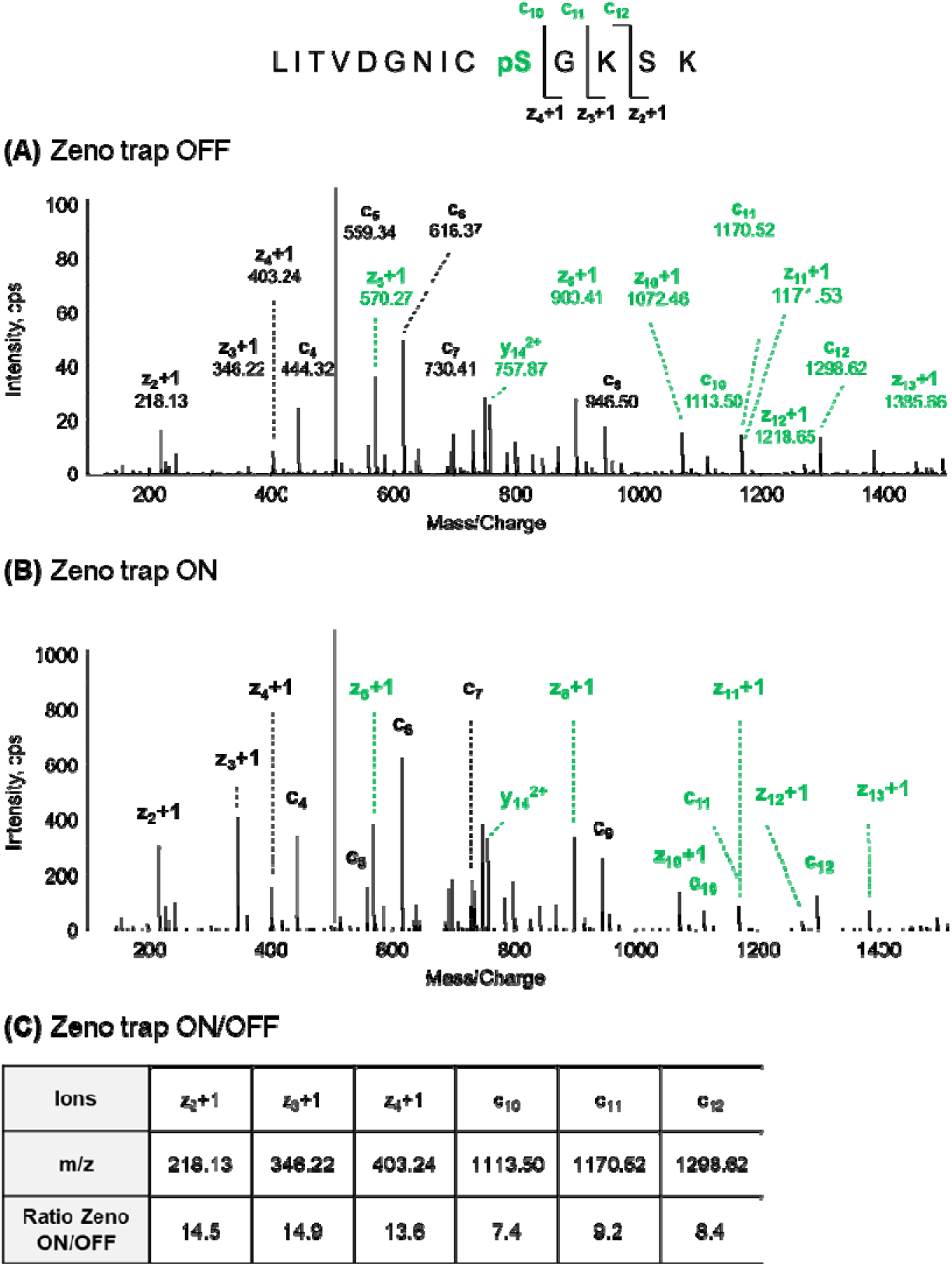
Increased EAD MS/MS sensitivity for phosphorylation analysis with Zeno trap technology. *EAD MS/MS spectra of the phosphorylated LITVDGNIC**^56^pS**GKSK precursor ion (m/z 505.58, z = 3+) **(A)** without and **(B)** with Zeno trap activated are shown. **(C)** LITVDGNIC**^56^pS**GKSK peptide was injected at 4 pg on-column for EAD PRM analysis with and without using the Zeno trap. Chromatographic peaks were extracted, and sensitivity changes with the Zeno trap on and off were determined. EAD KE = 2 eV was applied*.

Our data indicated that Zeno trap activation significantly increased EAD MS/MS spectral quality and sensitivity for all fragment ions. Combining EAD with Zeno trap activation strengthens the confidence for PTM site localization and isomer differentiation and is an asset to qualitatively and quantitatively analyze very low-abundance peptide ions and detect PTM site-specific “intact” fragment ions.

### Accurate quantification of PTM with targeted EAD PRM assays

To explore the quantitative performances of Zeno EAD PRM, four ranges of synthetic isomer modified peptides spiked at five different ratios were analyzed with Zeno trap activated and are detailed below.

#### Quantification accuracy

To first assess the relative quantification accuracy of Zeno EAD PRM vs Zeno CID PRM, we analyzed the doubly malonylated-acetylated peptide isoforms from human histone H2B type 1-C/E/F/G/I (P62807), PEPA**^6^Kmal**SAPAP**^12^Kac**K with a malonyl-lysine and acetyl-lysine at positions K-6 and K12, and PEPA**^6^Kac**SAPAP**^12^Kmal**K with an acetyl-lysine and malonyl-lysine at positions K-6 and K12. These isomers are hereafter referred to as M1 and M2, respectively, and were mixed at the following ratios of M1/M2: 0:4, 1:3, 2:2, 3:1, and 4:0. Each sample was analyzed by EAD PRM and CID PRM, with Zeno trap activated, and XICs of the site-specific z_7_+1/y_7_ (EAD/CID) ion were used for quantification with Skyline. Specifically, the differentiating z_7_+1 ion was used for relative quantification as it contains the acetyl group for M1 (z_7_+1: m/z 724.41) and the malonyl group for M2 (z_7_+1: m/z 768.40). **Figure 8A** displays the theoretical and EAD/CID experimental ratios for the different peptide isomers. Zeno EAD PRM estimated the five ratios very close to the expected values. On the contrary, results obtained for Zeno CID PRM showed that z_7_+1 ion with the acetyl group (represented in pink on **Figure 8A**) was primarily quantified, which reflects the abundant neutral losses due to CID fragmentation. Indeed, neutral loss of CO_2_ (-44 m/z) from the malonyl group results in an acetyl group on the lysine residue (**Figure 2D**). Consequently, the z_7_+1 ion from M2 was detected at m/z 724.41 as for z_7_+1 ion with the acetyl group from M1, but not at m/z 768.40 as with the malonyl group.

**Figure 8.**
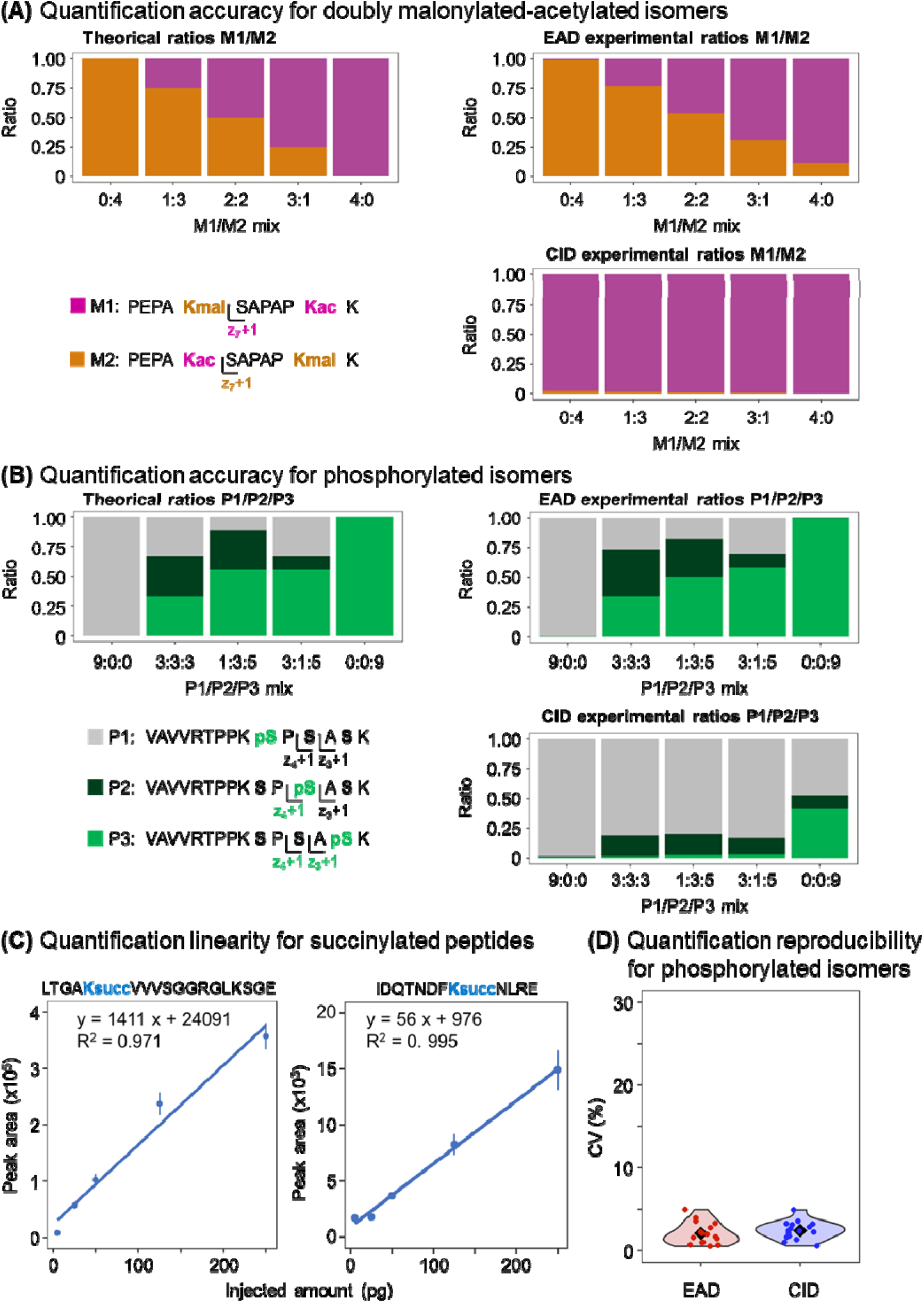
High quantification accuracy, linearity and reproducibility for PTM analysis by Zeno EAD PRM. (A-B) *Quantification accuracy was assessed by mixing modified peptide isomers at different ratios. (A) The doubly malonylated-acetylated peptide isoforms PEPA**^6^Kmal**SAPAP**^12^Kac**K (M1, in pink) and PEPA**^6^Kac**SAPAP**^12^Kmal**K (M2, in brown) (precursor ions at m/z 674.86, z = 2+), from human histone H2B type 1-C/E/F/G/I (P62807), were mixed at the following ratios of M1/M2: 0:4, 1:3, 2:2, 3:1 and 4:0, and analyzed by EAD PRM (KE = 2 eV) and CID PRM with Zeno trap activated. The chromatographic peak of the differentiating z_7_+1 or y_7_ ion was extracted for both peptides, and experimental M1/M2 ratios were determined. Zeno EAD PRM estimated the ratios very closely to the expected values, unlike Zeno CID PRM. Indeed, neutral losses from the labile malonyl group with CID fragmentation generate ions with the same m/z as acetylated ions. (B) The phosphorylated peptide isoforms VAVVRTPPK*^235^***pS****PSASK (P1, in grey), VAVVRTPPKSP*^237^***pS****ASK (P2, in dark green), and VAVVRTPPKSPSA*^239^***pS****K (P3, in green) (precursor ions at m/z 535.29, z = 3+), from mouse tau (P10637), were mixed at the following ratios of P1/P2/P3: 9:0:0, 3:3:3, 1:3:5, 3:1:5 and 0:0:9, and analyzed by EAD PRM (KE = 2 eV) and CID PRM with Zeno trap activated. The chromatographic peak of the differentiating z_4_+1/y_4_ ion for P1 and z_3_+1/y_3_ ion for P3 was extracted, and experimental P1/P2/P3 ratios were determined (P2 proportion was calculated as [1 – P1 proportion (from z_4_+1/y_4_) – P3 proportion (from z_3_+1/y_3_)]). Zeno EAD PRM estimated the ratios very closely to the expected values, contrary to Zeno CID PRM because of neutral losses of the phosphate group (-98 m/z) from phosphorylated serine residues. **(C)** Quantification linearity was assessed by mixing eight succinylated peptides, including LTGA*^216^***Ksucc****VVVSGGRGLKSGE (precursor ion at m/z 607.68, z = 3+) from mouse mitochondrial electron transfer flavoprotein subunit alpha (Q99LC5) and IDQTNDF**^76^Ksucc**NLRE (precursor ions at m/z 534.93, z = 3+) from mouse mitochondrial isovaleryl-CoA dehydrogenase, and spiking them at 5, 25, 50, 125 and 250 pg (on-column) in a constant medium complexity background. For each spiked sample, four injection replicates were collected by EAD PRM (KE = 7 eV) with Zeno trap activated. The chromatographic peak of PTM specific ions were extracted and used for quantification. Dots represent the mean value, error bars the standard deviation, and the linear regression is displayed. **(D)** Violin plots showing the coefficient of variation (CV) calculated on fragment ion peak area for the phosphorylated peptides P1, P2 and P3 analyzed in triplicate by EAD PRM (KE = 2eV) and CID PRM with Zeno trap activated. Peak area values of z_3_+1/y_3_ and z_4_+1/y_4_ ions obtained for the five previously described P1/P2/P3 mix samples were used (N = 16). The black diamond represents the mean*.

In addition, we analyzed the phosphorylated peptide isoforms from mouse tau (P10637), VAVVRTPPK^235^**pS**PSASK, VAVVRTPPKSP^237^**pS**ASK, and VAVVRTPPKSPSA^239^**pS**K, with a phospho-serine at positions S-235, S-237 and S-239, and hereafter referred to as P1, P2, and P3, respectively. These isomers were mixed at the following ratios of P1/P2/P3: 9:0:0, 3:3:3, 1:3:5, 3:1:5, and 0:0:9. Each sample was acquired in EAD PRM and CID PRM with Zeno trap activated, and then processed in Skyline. P1 was quantified using the differentiating z_4_+1 ion (m/z 376.20)/y_4_ ion (m/z 392.21) that does not contain phosphoryl groups, P3 using the differentiating z_3_+1 ion (m/z 369.13)/y_3_ ion (m/z 385.15) with one phosphoryl group, and P2 was estimated as the remaining proportion as no specific ion can be obtained. Similar results as for the malonylated-acetylated isomers were obtained with the phosphorylated isomers (**Figure 8B**): EAD estimated the different ratios with high accuracy. However, CID generated partial neutral loss of the phosphate group (-98 m/z) from phosphorylated serine residues, as observed for z_3_+1 ion from P3 (represented in green on **Figure 8B**), which consequently underestimated the abundance of z_3_+1 ion from P3 (one phosphoryl group) primarily toward overestimating z_4_+1 ion from P1 (no phosphoryl group). It is worth noting that CID quantification was partially recovered when monitoring both the intact ions and ions with neutral loss (**Figure S4**); however, this approach relies on “indirect” evidence and significantly complicates the data analysis. Altogether, these results demonstrate that, by preserving the labile PTM group, Zeno EAD PRM achieved very accurate quantification performances of modified peptides and, more strikingly, PTM peptide isomers.

#### Quantification linearity

To assess the linearity of EAD PRM, a mixture of eight synthetic non-tryptic succinylated peptides was prepared with various ratios, namely 2.5, 12.5, 25, 62.5 and 125 pg/μL, in a constant medium complex background. Each sample was acquired in quadruplicate in EAD PRM, with Zeno trap activated, and analyzed in Skyline using differentiating PTM site-specific ions. The mixture of succinylated peptides included the two isoforms from mouse mitochondrial electron transfer flavoprotein subunit alpha (Q99LC5), LTGA^216^**Ksucc**VVVSGGRGLKSGE and LTGAKVVVSGGRGL^226^**Ksucc**SGE, with a succinyl-lysine at positions K-216 and K-226. Near complete c and z+1 fragment ion series were obtained with EAD, and all differentiating ions (*i.e.,* c_5_ to c_14_ and z_4_+1 to z_13_+1 ions) were clearly detected on the EAD MS/MS spectra (ions containing the succinyl group are represented in blue, and additional differentiating ions in black) (**Figure S5**). It is worth noting that the sequence coverage was poorer with CID fragmentation as partial ion series were observed on the CID MS/MS spectra: succinyl-containing ions were under-represented for the two isomer peptides. Indeed, only b_7_ ion was clearly detected for LTGA^216^**Ksucc**VVVSGGRGLKSGE, and y_4_ and y_8_ to y_11_ ions for LTGAKVVVSGGRGL^226^**Ksucc**SGE (**Figure S6**). Quantification was performed in Skyline by extracting the chromatographic peaks of differentiating PTM site-specific ions (*i.e.*, z_4_+1, z_5_+1, z_7_+1, z_9_+1, z_10_+1, c_8_, c_9_, c_11_, c_12_ and c_13_) (**Figure S5**). The linear regression R^2^ equaled 0.971 for LTGA^216^**Ksucc**VVVSGGRGLKSGE (**Figure 8C**) and 0.946 for LTGAKVVVSGGRGL^226^**Ksucc**SGE (**Figure S7A**). The R^2^ of the linear regression reached 0.995 for the succinylated peptide IDQTNDF**^76^Ksucc**NLRE from mouse mitochondrial isovaleryl-CoA dehydrogenase (Q9JHI5) with a succinyl-lysine at position K-76 (**Figure 8C**). Quantification linearity was investigated for five more succinylated peptides, and R^2^ ranged between 0.978 and 0.998 (**Figure S7B-F**) with a median R^2^ value of 0.993 for the eight succinylated peptides. Thus, Zeno EAD PRM achieved very good linearity performance on modified peptides.

#### Quantification reproducibility

Reproducibility is a crucial aspect for qualitative and quantitative investigations by MS-based proteomics. While targeted MS is well recognized as a highly reproducible strategy ^34^, reproducibility of targeted PRM applying EAD or CID fragmentation on the recent QqTOF mass spectrometer for the analysis of peptides with labile PTM group needs to be demonstrated. To do so, the coefficients of variation (CVs) of z_3_+1/y_3_ and z_4_+1/y_4_ (EAD/CID) ion peak areas were determined for the phosphorylated isomers VAVVRTPPK^235^**pS**PSASK (P3), VAVVRTPPKSP^237^**pS**ASK (P4), and VAVVRTPPKSPSA^239^**pS**K (P5) in the five described P6/P7/P8 mix samples injected in triplicate (**Figure 8D**). On average, CVs equaled 2.1% and 2.4% for EAD PRM and CID PRM, respectively. This highlights a high degree of reproducibility achieved by both EAD PRM and CID PRM techniques conducted on the QqTOF instrument.

## CONCLUSIONS

EAD is a recently introduced electron-based fragmentation technology for LC-MS/MS analysis. More specifically, for PTM analysis by tandem mass spectrometry, EAD with a low kinetic energy electron beam prevents neutral losses from labile PTM groups (e.g., malonylation and phosphorylation) and provides comprehensive c and radical z+1 fragmentation series for both tryptic and non-tryptic peptides with strong PTM-site localizing ions. Thus, EAD fragment ion spectra provide the single amino-acid resolution required for isomer differentiation and respective PTM site identification. More importantly, EAD kinetic energy (KE) can be optimized and fine-tuned for specific PTMs. This is a powerful feature with respect to electron-based activation methods. Indeed, tunable KE enables the generation of intact ions with the PTM group, providing direct and highly confident evidence of the PTM sites, as well as peptide sequence information. Here, optimal KEs ranged between 2 eV and 7 eV for the different modification groups and peptide sequences investigated, while higher KE values induced partial neutral losses of the labile malonyl and phosphoryl groups. Interestingly, high EAD KE can be employed to promote characteristic neutral losses from the modification groups, that can yield additional evidence for PTM identification and localization.

EAD has been implemented on a recent QqTOF mass spectrometer and the EAD cell is located directly in front of the Q2 quadrupole collision cell in the ion path. This MS system is also equipped with a Zeno trap device located directly after the collision cell, which when activated yields almost 100% duty cycle for delivery of MS/MS fragment ions into the TOF accelerator region and, consequently, an increase in MS/MS signal intensity and sensitivity. In this study, activating the Zeno trap achieved an average 6– 11-fold gain for EAD PRM analyses. Thus, combining EAD with Zeno trap activation on the highly sensitive and fast-scanning QqTOF system confidently and accurately identified peptides and located the PTM sites, even at low levels.

Preliminary quantitative analyses on various PTM types and isomers demonstrated accurate, linear, and reproducible performances for quantifying EAD fragment ions by targeted PRM. Indeed, the ability of EAD to maintain labile groups intact is of great benefit for achieving high quantification performance and accurate site-specific ion quantification, while neutral losses induced by CID might lead to inaccurate relative quantification, specifically when studying modified peptide isomers.

Various strategies are available for analyzing Zeno EAD PRM data. We showed that collected data can be processed using the SCIEX OS software for data visualization and XIC extraction and quantification, as well as the open-source, platform-independent, and vendor agnostic Skyline software platform. Skyline computes c ion, z+1 radical ion (z^•^ ion) and z+2 ion (z’ ion) types from EAD, ECD, and ETD data that can be visualized and used for precise MS/MS quantification.

In conclusion, our work highlights novel workflows and approaches for PTM identification, site localization, and quantification. Here we applied EAD to acylation (malonylation, acetylation, and succinylation) and phosphorylation, but these methods can be used to investigate other types of peptide and protein modifications. Obtaining confident and precise information on PTM identification, site localization, and quantification is essential to understand how proteins are affected by modifications and, thus, to gain deeper insights into biological processes and disease mechanisms.

## AUTHOR INFORMATION

### Corresponding Author

* Birgit Schilling – Buck Institute for Research on Aging, 8001 Redwood Blvd, Novato, CA 94945, USA; Phone: +1 (415) 209-2079;

### Author Contributions

C.L.H. and B.S. designed the experiments. J.B., C.L.H., J.C., and B.S. performed the experiments. J.B., C.L.H., R.C., J.P.R., A.A., B.ML, and B.S. performed data analysis. J.B., C.L.H., and B.S. wrote the manuscript. All authors have approved the final version of the manuscript.

### Notes

C.L.H., J.C., and A.A. are employees of SCIEX.

## Supporting information

meta data

## ACKNOWLEDGMENT

The authors acknowledge the generous support from SCIEX for the Waters M-Class HPLC and ZenoTOF 7600 mass spectrometer at the Buck Institute.

