## Supplementary Figure for "Localization and Quantification of Post-Translational Modifications of Proteins Using Electron Activated Dissociation Fragmentation on a Fast-Acquisition Time-of-Flight Mass Spectrometer"

### Supplementary Figure S1

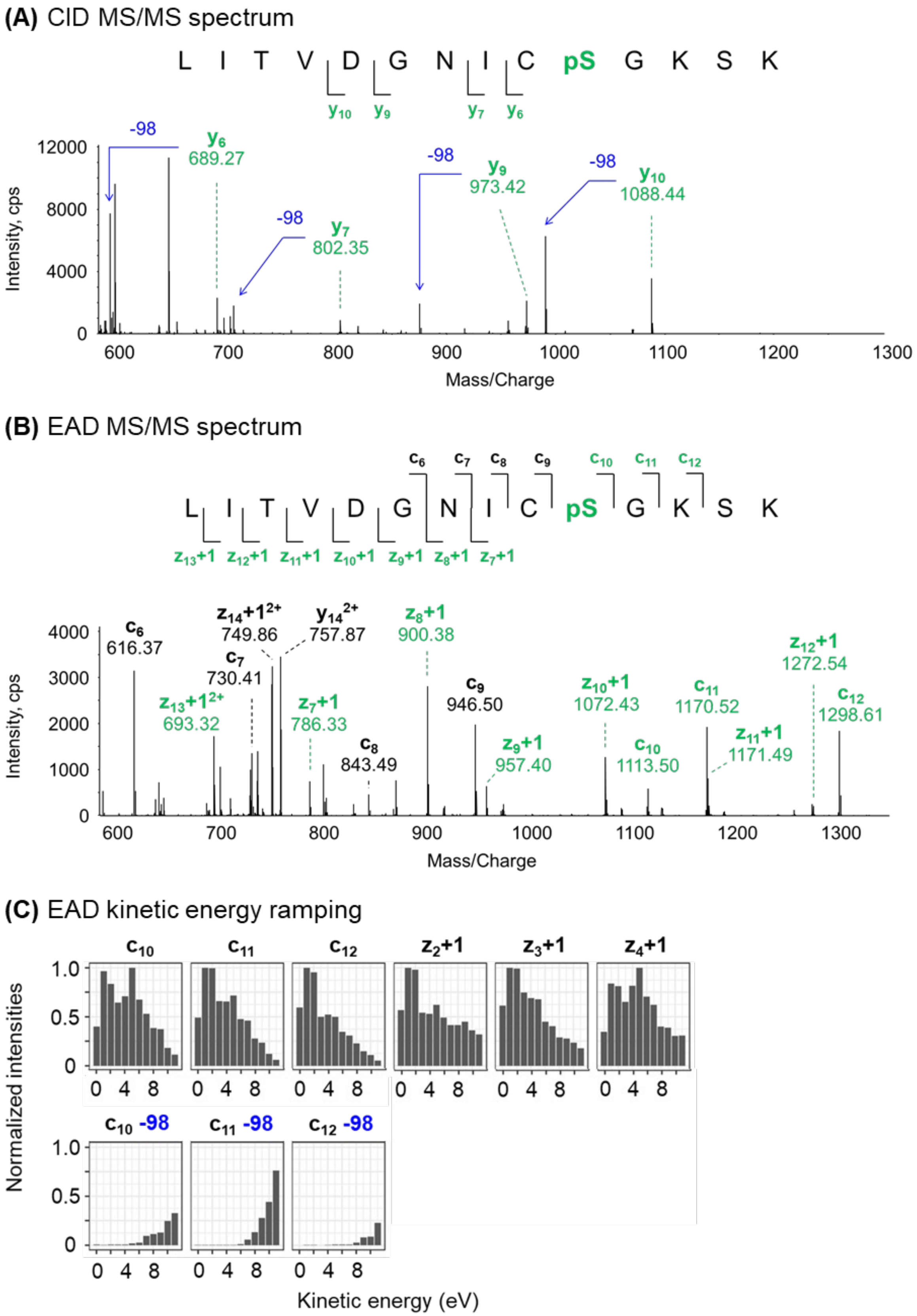

**Supplementary Figure S1. Preservation of the labile phosphoryl group of LITVDGNICpSGKSK at low EAD kinetic energy.** The phosphorylated peptide LITVDGNIC[56]**pS**GKSK (precursor ion at m/z 505.58, z = 3+) from the bovine mitochondrial NDUFA10 subunit of Complex I (P34942) was analyzed using **(A)** Zeno CID PRM and **(B)** Zeno EAD PRM (KE = 2 eV). Site-specific fragment ions containing the phosphoryl group are indicated in green. Neutral losses (-98 m/z) were observed in CID. **(C)** The EAD kinetic energy was ramped from 0 eV to 11 eV. Chromatographic peaks were extracted for nine fragment ions. For each intact fragment ion (first line), peak area values were normalized to the respective highest area value. For the fragment ions with a neutral loss (second line), area values were normalized to the value of the intact ion counterpart, respective of the KE. Neutral losses (-98 m/z) were observed for higher KEs. Displayed MS/MS were zoomed over an m/z range of 600-1,300, and the full scan EAD MS/MS is shown in **Figure 4**.

#### Supplementary Figure S2

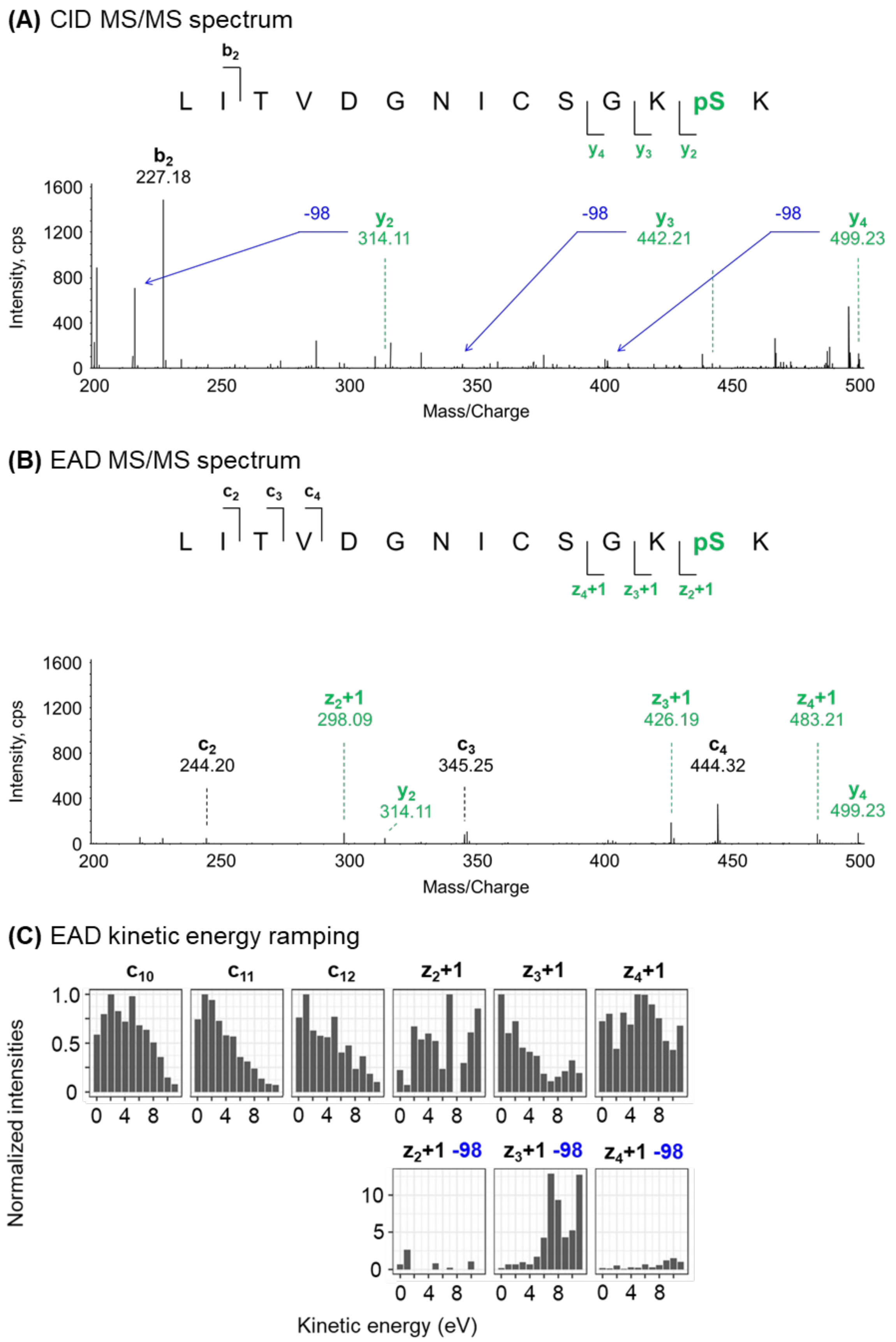

**Supplementary Figure S2. Preservation of the labile phosphoryl group of LITVDGNICSGKpSK at low EAD kinetic energy.** The phosphorylated peptide LITVDGNICSGK[59]**pS**K (precursor ion at m/z 505.58, z = 3+) from the bovine mitochondrial NDUFA10 subunit of Complex I (P34942) was analyzed using **(A)** Zeno CID PRM and **(B)** Zeno EAD PRM (KE = 2 eV). Site-specific fragment ions containing the phosphoryl group are indicated in green. Neutral losses (-98 m/z) were observed in CID. **(C)** The EAD kinetic energy was ramped from 0 to 11 eV. Chromatographic peaks were extracted for nine fragment ions. For each intact fragment ion (first line), peak area values were normalized to the respective highest area value. For the fragment ions with a neutral loss (second line), area values were normalized to the value of the intact ion counterpart, respective of the KE. Neutral losses (-98 m/z) were observed for higher KEs. Displayed MS/MS were zoomed over an m/z range of 200-500, and the full scan EAD MS/MS is shown in **Figure 4**.

### Supplementary Figure S3

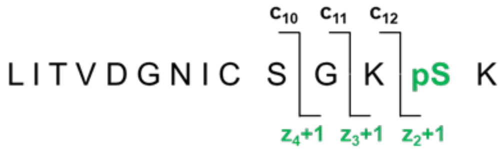


**(A)** Zeno trap OFF

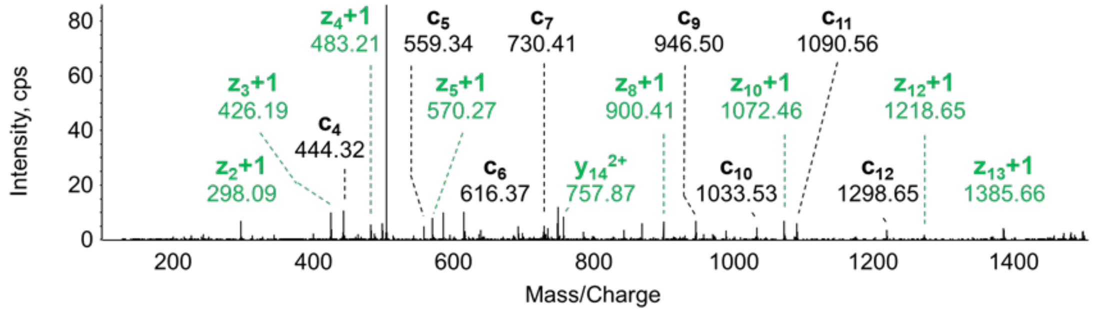


**(B)** Zeno trap ON

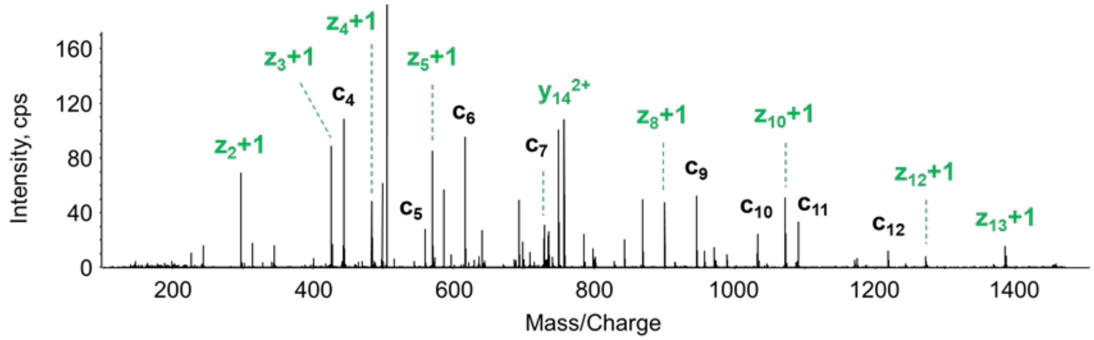


**(C)** Zeno trap ON/OFF

| **Ions** | $z_2$+1 | $z_3$+1 | $z_4$+1 | $c_{10}$ | $c_{11}$ | $c_{12}$ |
|---|---|---|---|---|---|---|
| **m/z** | 298.09 | 426.19 | 483.21 | 1033.53 | 1090.56 | 1218.65 |
| **Ratio Zeno ON/OFF** | 10.4 | 12.7 | 19.9 | 8.8 | 9.9 | 6.8 |

**Supplementary Figure S3. Another example showing the gain of sensitivity provided by Zeno trap activated for EAD phosphorylation analysis.** EAD MS/MS spectra of the phosphorylated LITVDGNICSGK$^{59}$**pS**K precursor ion (m/z 505.58, z = 3+) **(A)** without and **(B)** with Zeno trap activated. The y-axis scale is different for the spectra. **(C)** LITVDGNICSGK$^{59}$**pS**K peptide was injected at 4-pg on-column for EAD PRM analysis with and without using the Zeno trap. Chromatographic peaks were extracted, and sensitivity changes between Zeno trap on and off were determined. EAD KE = 2 eV was applied.

### Supplementary Figure S4

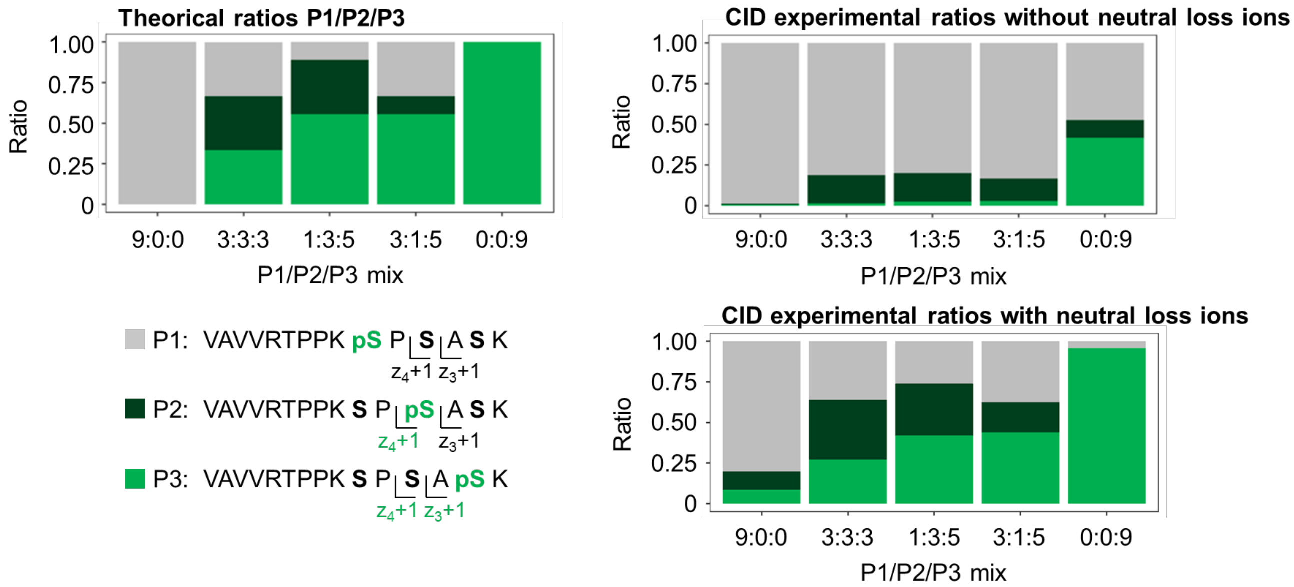

**Supplementary Figure S4. Quantification accuracy for serine phosphorylation analysis by Zeno CID PRM.** Quantification accuracy was assessed by mixing the phosphorylated peptide isoforms VAVVRTPPK$^{235}$**pS**PSASK (P1, in grey), VAVVRTPPKSP$^{237}$**pS**ASK (P2, in dark green), and VAVVRTPPKSPSA$^{239}$**pS**K (P3, in green) (precursor ions at m/z 535.29, z = 3+), from mouse tau (P10637), at the following ratios of P1/P2/P3: 9:0:0, 3:3:3, 1:3:5, 3:1:5 and 0:0:9. Samples were analyzed by CID PRM with Zeno trap activated. The chromatographic peaks of the differentiating phosphoryl group-containing $y_4$ ion for P1 and $y_3$ ion for P3 as well as the corresponding ions resulting from neutral loss, $y_4$-98 ion for P1 and $y_3$-98 ion for P3, were extracted. Then, areas of [$y_4$ + $y_4$-98] and [$y_3$ + $y_3$-98] were calculated to estimate P1 and P3 proportion, respectively. Finally, experimental P1/P2/P3 ratios were determined (P2 proportion was calculated as [1 – P1 proportion – P3 proportion]). The proportions of P1, P2, and P3 corrected with the neutral loss ions appeared closer to the expected ratios than when the neutral loss ions are not included.

### Supplementary Figure S5

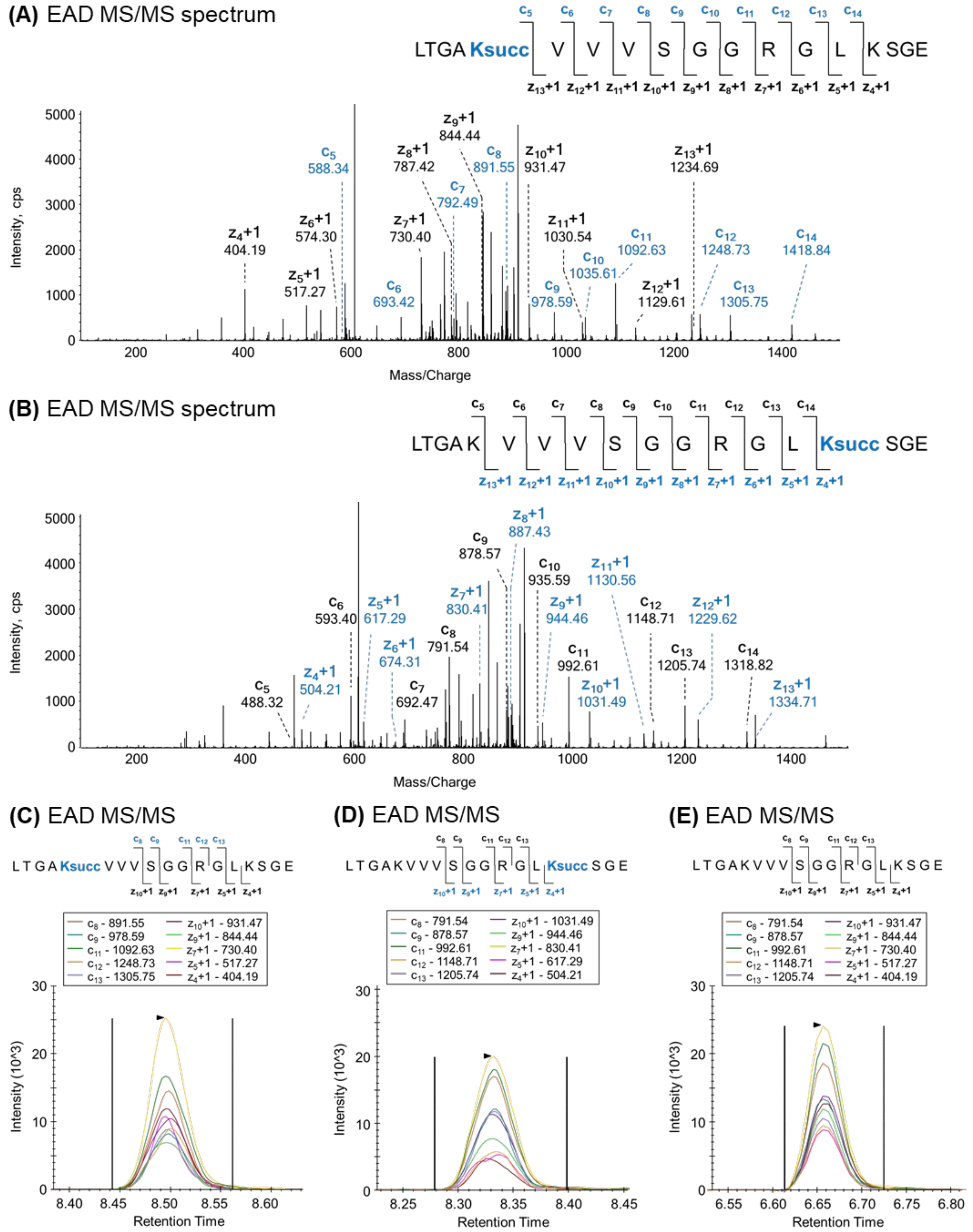

**Supplementary Figure S5. Confident differentiation of succinylated peptide isomers using EAD.** Zeno EAD MS/MS spectra of the succinylated **(A)** LTGA$^{216}$**Ksucc**VVVSGGRGLKSGE and **(B)** LTGAKVVVSGGRGL$^{226}$**Ksucc**SGE isomer precursor ions (m/z 607.68, z = 3+) from mouse mitochondrial electron transfer flavoprotein subunit alpha (Q99LC5) are shown for 250-pg on-column injections. EAD KE = 7 eV was applied. Site-specific fragment ions containing the succinyl group are indicated in blue, and additional differentiating ions are in black. Differentiating ions of succinylated isomers **(C)** LTGA$^{216}$**Ksucc**VVVSGGRGLKSGE and **(D)** LTGAKVVVSGGRGL$^{226}$**Ksucc**SGE and **(E)** their unmodified counterpart LTGAKVVVSGGRGLKSGE analyzed in EAD were extracted in Skyline for quantification.

### Supplementary Figure S6

**(A)** CID MS/MS spectrum

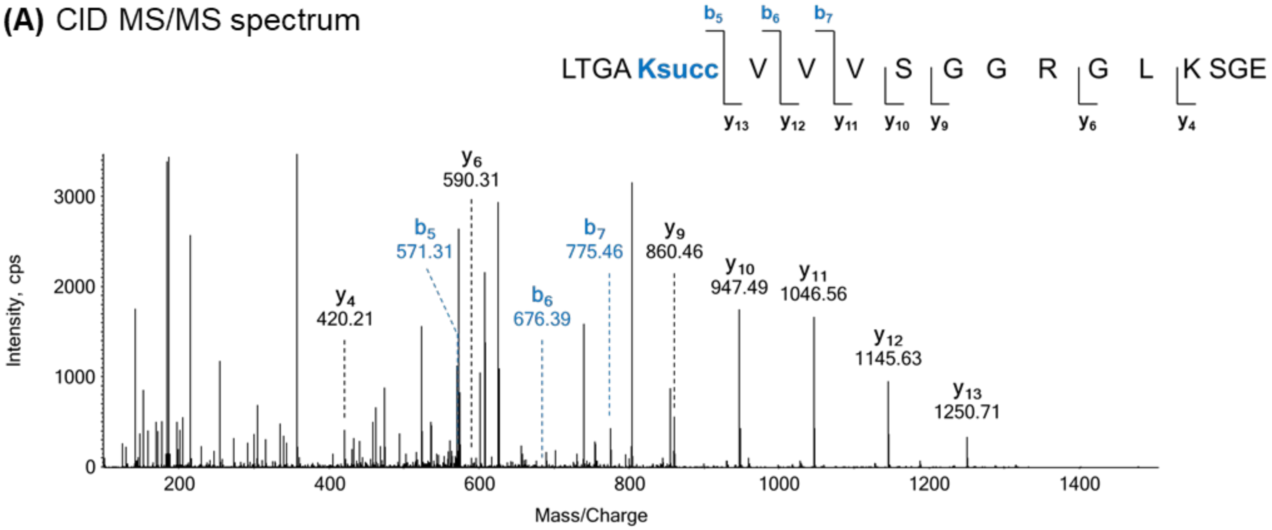


**(B)** CID MS/MS spectrum

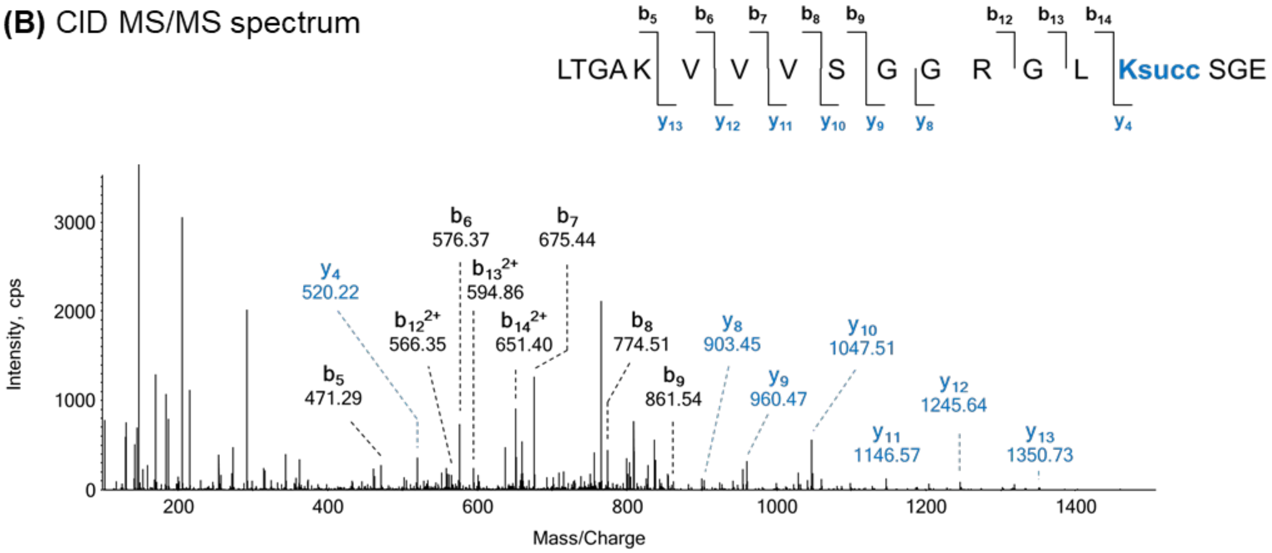

**Supplementary Figure S6. Incomplete fragmentation ion series of succinylated peptide isomers using CID.** Zeno CID MS/MS spectra of the succinylated **(A)** LTGA[216]**Ksucc**VVVSGGRGLKSGE and **(B)** LTGAKVVVSGGRGL[226]**Ksucc**SGE isomer precursor ions (m/z 607.68, z = 3+) from mouse mitochondrial electron transfer flavoprotein subunit alpha (Q99LC5) are shown for 50-pg on-column injections. Site-specific fragment ions containing the succinyl group are indicated in blue, and additional differentiating ions are in black.

#### Supplementary Figure S7

**(A)** Quantification linearity for succinylated LTGA**K**VVVSGGRGL**Ksucc**SGE

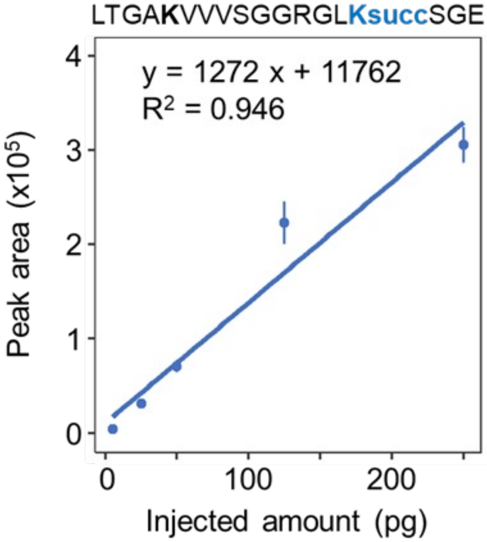


**(B)** Quantification linearity for succinylated GA**Ksucc**YVSHGATGKGNDQVRFE

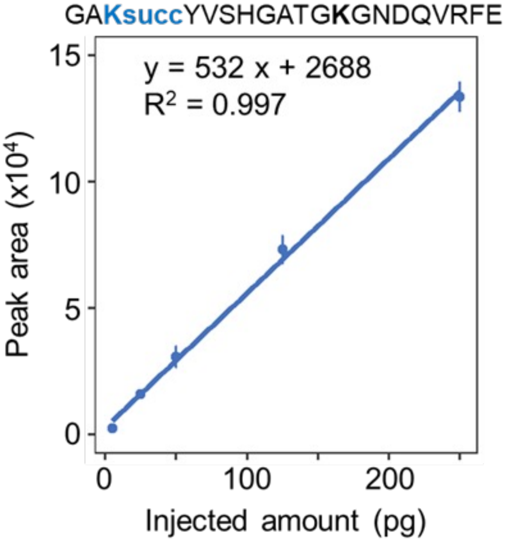


**(C)** Quantification linearity for succinylated LQHHV**Ksucc**SVTAPYKYPRKVE

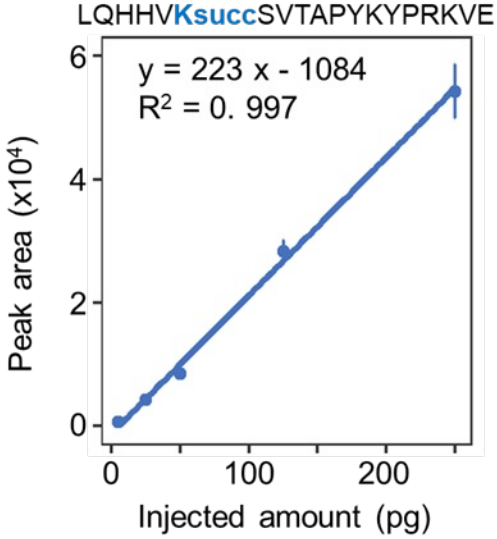


**(D)** Quantification linearity for succinylated SLKRMAK**Ksucc**KFTENPKAGDE

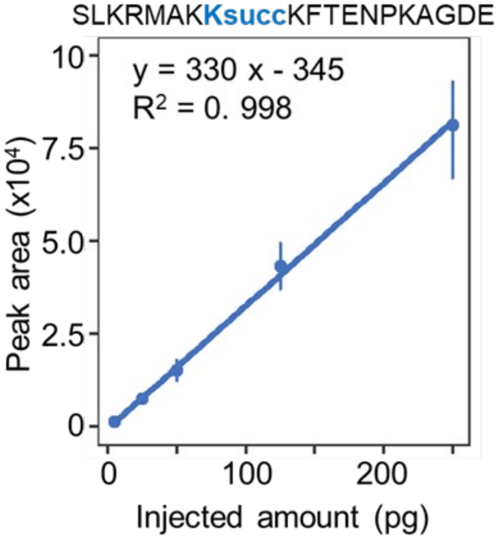

#### Supplementary Figure S7

**(E)** Quantification linearity for succinylated VIKTPMTSQ**Ksucc**TFE

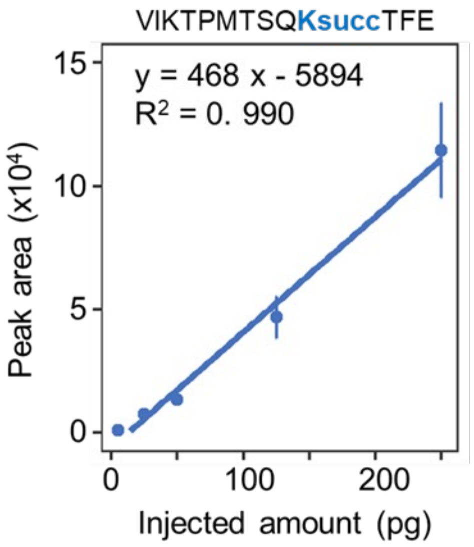


**(F)** Quantification linearity for succinylated VNPTLG**Ksucc**TAEE

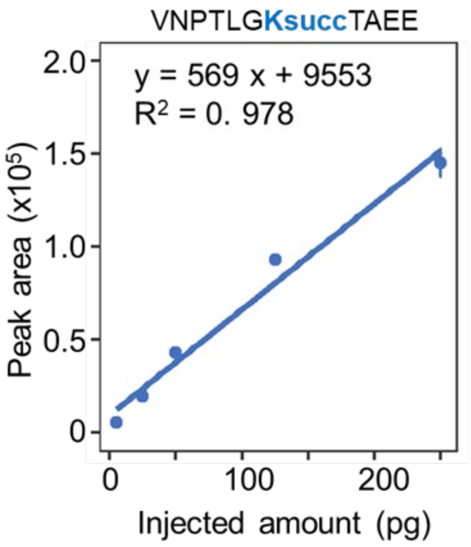

**Supplementary Figure S7. Quantification linearity for lysine succinylation analysis by Zeno EAD PRM.** Quantification linearity was assessed by mixing succinylated peptides at five various ratios (5, 25, 50, 125, and 250 pg on-column) in a constant medium complexity background. For each spiked sample, four injection replicates were collected by EAD PRM (KE = 7 eV) with Zeno trap activated. The chromatographic peak of differentiating ions was extracted in Skyline and used for quantification. Dots represent the mean value, error bars the standard deviation, and the linear regression is displayed. **(A)** Succinylated peptide LTGAKVVVSGGRGL$^{226}$**Ksucc**SGE (precursor ions at m/z 607.68, z = 3+) from mouse mitochondrial electron transfer flavoprotein subunit alpha (Q99LC5). **(B)** Succinylated peptide GA$^{112}$**Ksucc**YVSHGATGKGNDQVRFE (precursor ions at m/z 557.52, z = 4+) from mouse argininosuccinate synthase (P16460). **(C)** Succinylated peptide LQHHV$^{534}$**Ksucc**SVTAPYKYPRKVE (precursor ions at m/z 597.33, z = 4+) from mouse mitochondrial acyl-coenzyme A synthetase ACSM1 (Q91VA0). **(D)** Succinylated peptide SLKRMAK$^{80}$**Ksucc**KFTENPKAGDE (precursor ions at m/z 572.80, z = 4+) from mouse mitochondrial hydroxyacyl-coenzyme A dehydrogenase (Q61425). **(E)** Succinylated peptide VIKTPMTSQ$^{192}$**Ksucc**TFE (precursor ions at m/z 808.42, z = 2+) from mouse mitochondrial hydroxyacyl-coenzyme A dehydrogenase (Q61425). **(F)** Succinylated peptide VNPTLG$^{284}$**Ksucc**TAEE (precursor ions at m/z 632.82, z = 2+) from mouse arginase-1 (Q61176).
